# Rapid Proteomic Screen of CRISPR Experiment Outcome by Data Independent Acquisition Mass Spectrometry: A Case Study for HMGN1

**DOI:** 10.1101/490763

**Authors:** Martin Mehnert, Wenxue Li, Chongde Wu, Barbora Salovska, Yansheng Liu

## Abstract

CRISPR-Cas gene editing holds substantial promise in many biomedical disciplines and basic research. Due to the important functional implications of non-histone chromosomal protein HMG-14 (HMGN1) in regulating chromatin structure and tumor immunity, we performed gene knockout of HMGN1 by CRISPR in cancer cells and studied the following proteomic regulation events. In particular, we utilized DIA mass spectrometry (DIA-MS) and reproducibly measured more than 6200 proteins (protein-FDR 1%) and more than 82,000 peptide precursors in the single MS shots of two hours. HMGN1 protein deletion was confidently verified by DIA-MS in all of the clone- and dish- replicates following CRISPR. Statistical analysis revealed 147 proteins changed their expressions significantly after HMGN1 knockout. Functional annotation and enrichment analysis indicate the deletion of HMGN1 induces the histone inactivation, various stress pathways, remodeling of extracellular proteomes, cell proliferation, as well as immune regulation processes such as complement and coagulation cascade and interferon alpha/ gamma response in cancer cells. These results shed new lights on the cellular functions of HMGN1. We suggest that DIA-MS can be reliably used as a rapid, robust, and cost-effective proteomic screening tool to assess the outcome of the CRISPR experiments.

## Introduction

The clustered regularly interspaced short palindromic repeat (CRISPR) genome-editing system was demonstrated to be a revolutionary tool in many biological and biomedical fields, including studying gene functions, creating new cell and animal models, developing therapeutic agents, and potentially to treat diseases. CRISPR-Cas9 was adapted from a naturally occurring genome editing system in prokaryotic organisms that provides a form of acquired immunity [1]. In this system, CRISPR sequences are DNA fragments from viruses that have previously infected the prokaryote and are used to detect and destroy DNA from similar viruses during subsequent infections, whereas Cas9 (or ‘CRISPR-associated 9’) is an enzyme that cuts foreign DNA. Usually, the Cas9 nuclease (or other enzymes) can be delivered with a synthetic guide RNA (gRNA) into a cell so that the cell’s genome can be cut at a desired location [2–6]. Following DNA cleavage, researchers can use the eukaryotic cell’s own repair systems to either fuse broken chromosomal ends together via non-homologous end joining (NHEJ) or, in the presence of donor DNA, introduce exogenous sequence via homology-directed repair (HDR), for the purpose of gene knockout or knock in respectively [2–5].

A growing number of applications of CRISPR-Cas9 require a better understanding, characterization, and management of CRISPR efficacy and off-target effects. Many prediction tools and experimental approaches have been developed for these aims [7–10]. For example, Cas9 nickase activity was used to significantly reduce off-target activity and the application of paired guide RNAs (gRNAs) was shown to increase the efficiency of gene knockouts [11, 12]. Currently, the successful implementation of CRISPR experiment for a target gene or gene set is routinely validated by PCR using primers flanking the deleted region at the DNA level or by western blotting of the protein targets at the protein level. The next generation sequencing (NGS) techniques using e.g., amplicon-based kits have been successfully applied for screening CRISPR-Cas9 induced mutations, efficacy, and both on-target and off-target effects [13–16].

Due to the limitations of antibody such as cross-reactivity, the mass spectrometry (MS) based methods such as targeted proteomics [17, 18] have been widely adopted to detect and quantify proteins of interest. Over the last decade, the proteome has become precisely measurable. The acute state of the proteome—the *proteotype*—is a useful indicator of the cellular state and important for studying disease phenotype, development, pharmacological responses, and many other aspects in life science research [19, 20]. The last five years have witnessed significant technological improvements in MS-based proteomics especially in proteome coverage and measurement reproducibility of large sample cohorts. Particularly, data-independent acquisition (DIA) methods, such as SWATH mass spectrometry (SWATH-MS) combined with targeted data extraction has achieved unprecedented reproducibility, which are capable of generating quantitative matrices for thousands of proteins measured across multiple samples [20–22]. Recently, SWATH-MS was proven to be reproducible in a cross-lab study for >4000 proteins quantified in mammalian cells [23]. Moreover, the statistical strategies controlling the quality of protein identification and quantification in large-scale DIA-MS have matured [24, 25]. Because proteins catalyze and control directly most biological processes, and because *proteotype* characterization essentially bridges the genotypic variation and phenotype diversity, we propose that a sensitive, reproducible, robust, and rapid proteomic measurement enabled by e.g., DIA-MS would provide a general and powerful alternative approach to assess the outcome of a CRISPR experiment, to not only verify the successful knockout of the target gene (and its protein expression), but also to profile perturbed signaling pathways and quantitative phenotypes.

Herein, we focused on an example of a human protein, non-histone chromosomal protein HMG-14 (High Mobility Group Nucleosome Binding Domain 1, or HMGN1). Others and we have previously observed the HMGN1 mRNA and protein overexpression in Down Syndrome (DS, trisomy 21), which supports the hypothesis that HMGN1 could increase the chromatin accessibility and cause transcriptional dysregulation [26, 27]. Based on previous literatures, HMGN1 is associated to two major processes, transcriptional regulation and tumor immunity [28–36]. Despite of the emerging importance of HMGN1, there are currently no proteomic data sets available to understand the *proteotype* induced by its exclusive dosage imbalance in cells. In the present study we therefore analyzed the HMGN1 knockout cancer cells by DIA-MS to further understand the functional roles of HMGN1.

## Results

To specifically knock out HMGN1 by CRISPR, we transfected the human T-REx-HeLa CCL2 cells with two gRNAs that target the second and third exon of HMGN1, respectively. Single cells with positive GFP signal indicating Cas9 and gRNA expression were then sorted and expanded. To investigate the proteomic variation between these cell clones, three of them (named Clone A, B, and C hereafter) were verified by western blot to be negative with HMGN1 antibody staining, as compared to the wildtype control cells (**Figure 1**, **left panel**), and included in the study as *clone*- replicates. For each of A, B, C clones and the control, three independent cell cultures were included as *dish*- replicates. These 12 samples were randomized, processed, and measured by a two-hour DIA-MS on Orbitrap Lumos. Quantification was preformed based on high-resolute fragment signals (MS2 resolution, 30,000), followed by bioinformatic analysis (**Figure 1**).

**Figure 1.**
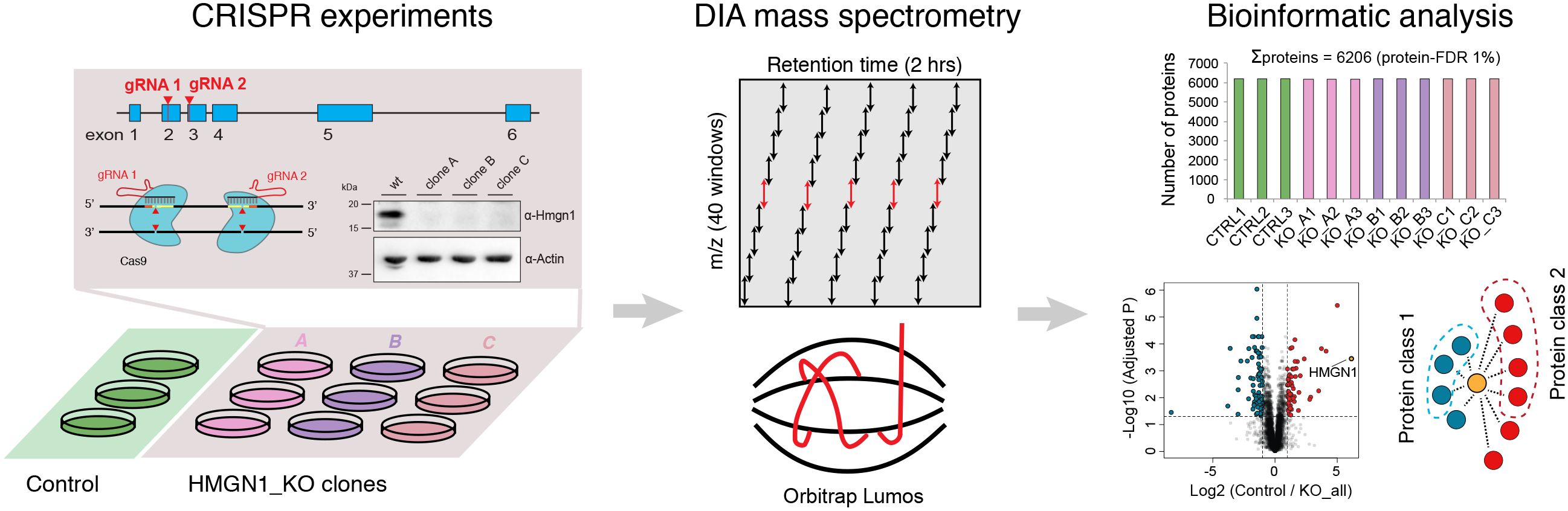
Experimental schema. Two gRNAs targeting the second and third exon of HMGN1 were used for the CRISPR/Cas9-mediated deletion. Three knockout clone- replicates and three dish- replicates per each clone and control cells were processed and measured by DIA proteomics, followed by bioinformatics.

To analyse the DIA-MS data, we used both spectrum-centric [37] (named directDIA in Spectronaut, a method extracting pseudo-spectrum directly from DIA data without the need of an assay library [38, 39]) and peptide-centric (based on SWATH assay library which contains mass spectrometric assays for 10,000 human proteins [40], named panHuman hereafter) approaches. Both approaches achieved substantial sensitivity when both peptide- and protein-FDR were strictly controlled at 1% (**Figure 2 A-B**) [25]. On average, directDIA identified and quantified 5,465 ± 7 Swiss-Prot proteins, assigned from 61,476 ± 430 unique peptides (~82,840 ± 685 peptide precursors) in the 12 samples, whereas panHuman library detected 5625 ± 27 proteins corresponding to 57,920 ± 1,601 unique peptides (~68,338 ± 2,092 peptide precursors). As expected, the majority of the proteins identified (i.e., 4,949 proteins) were shared between directDIA and panHuman (**Figure 2C**). Interestingly, despite directDIA quantified 2.83% less proteins, this library-free approach profiled 6.14% more peptides than panHuman library based analysis. This could be partially explained by the fact that the panHuman library was originally derived from TripleTOF platform [40], but still indicates the decent proteomic recovery, essentially at the peptide level, achieved by directDIA. Impressively, directDIA reaches 98.9% of data completeness in measuring all the precursors (**Supplementary Figure 1**). Taken together, we quantified 6,206 proteins in the combined results from the single measurements of two hours, which represents 50-60% of the total proteome expressed in a cancer cell line under a given condition [41].

**Figure 2.**
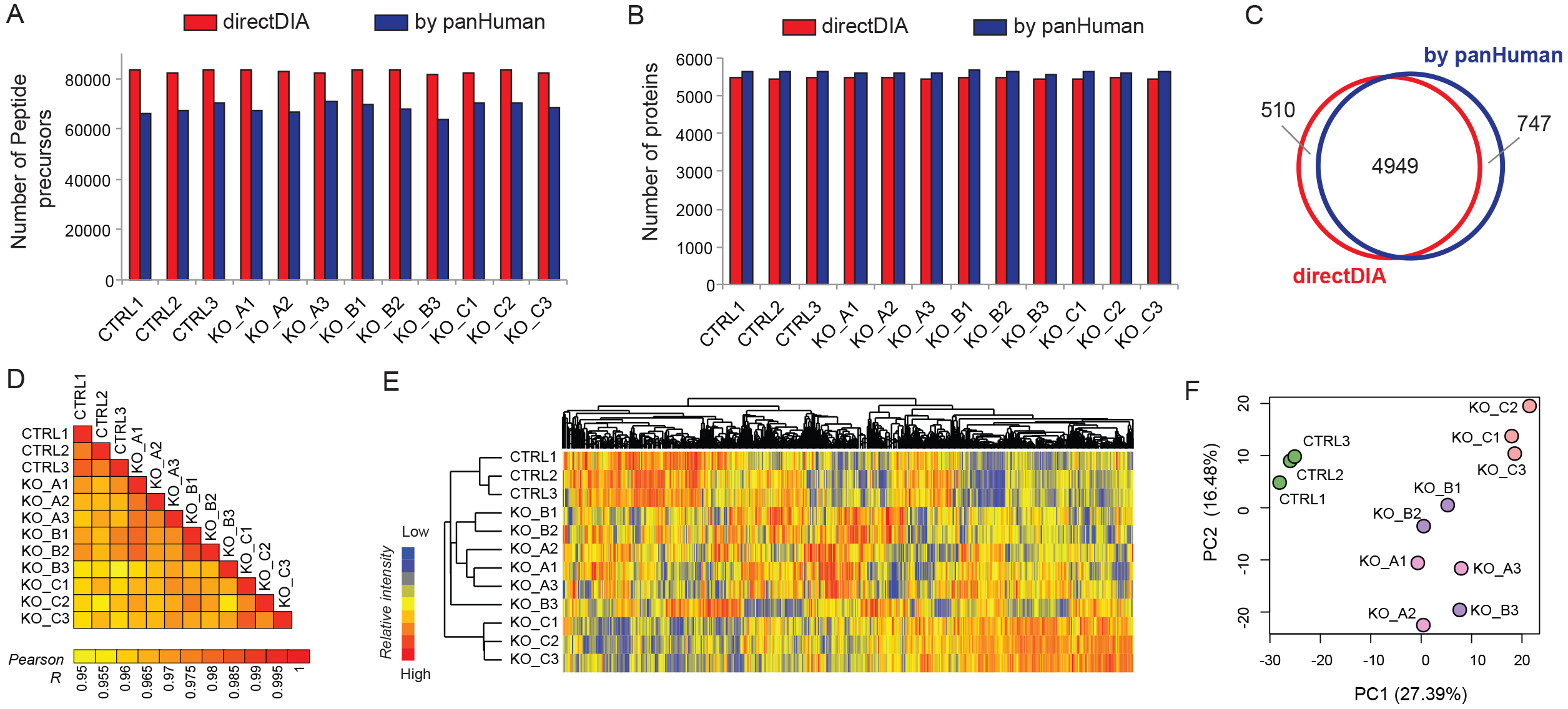
Identification and quantification results by DIA-MS. **(A)** Distribution of the number of peptide precursors identified in each sample. KO, knockout. **(B)** Distribution of non-redundant Swiss-Prot protein groups in each sample. **(C)** Venn diagram of protein identities identified by directDIA (library free) and panHuman library based approaches. **(D)** Sample-to-sample Pearson correlation coefficient R visualized. **(E)** Hierarchical clustering analysis (HCA) on median normalized, log2-transformed DIA-MS data. **(F)** Principal component analysis (PCA) between 12 samples. PC, principal component.

We next focused on analyzing the quantitative reproducibility and variation in both *clone*- and *dish*- replicates. Firstly, we found that 5487, 5277, 4361, and 5501 proteins of Control, Clone A, B, and C were quantified with a CV<20% in the three dish-replicates. The *sample-to-sample* Pearson correlations were fairly high throughout the whole experiment (**Figure 2D**), with e.g., average R=0.982 achieved between *dish*-replicates in Controls across five orders of magnitude of absolute DIA MS2 intensity (see **Supplementary Figure 2** for an example). Secondly, we performed both hierarchical clustering analysis (HCA) and principal component analysis (PCA) on the relative quantification results (**Figure 2 E-F**). In both plots, we found the three *dish*- replicates of Control and Clone C could emerge as individual clusters whereas Clone A and B *dish*-replicates were not separated from each other. The fact that Clone C showed deviation from to A and B could suggest a bigger clonal effect in C that was successfully revealed by DIA-MS (**Figure 2F**). In the meanwhile, Clone C has similar growth and proliferation rate to A and B (**Supplementary Figure 3**). Consistent to PCA result, the average CV for Clone B *dish*- replicates was 17.4%, which was higher than that of other clones (i.e, 11.41%, 13.21%, and 10.13% for Control, Clone A and C respectively, **Supplementary Figure 4**), suggesting a bigger *dish*-variation in B. Finally, all the 12 samples had an average CV of 21.56%, higher than that of each clones. Also, PCA plot indicates Control samples were separated from all knockout samples using the first principal component, suggesting the proteomic variation due to HMGN1 depletion was successfully captured in our whole data set. Overall, DIA-MS yielded excellent quantification reproducibility, providing an opportunity to faithfully reveal the *de facto dish*- and *clone*- variations, and to reveal HMGN1 dosage associated proteotype changes.

**Figure 3.**
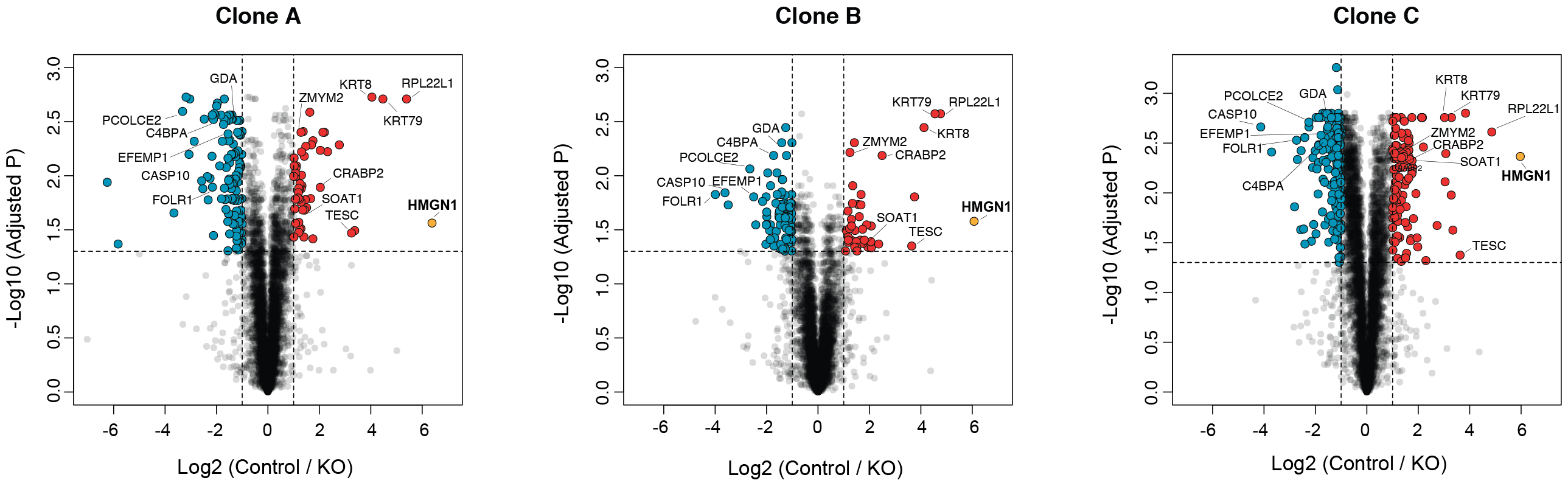
Volcano plots for the differential protein profiling between control and knockout clones using three dish- replicates. HMGN1 was highlighted by the yellow dot.

**Figure 4.**
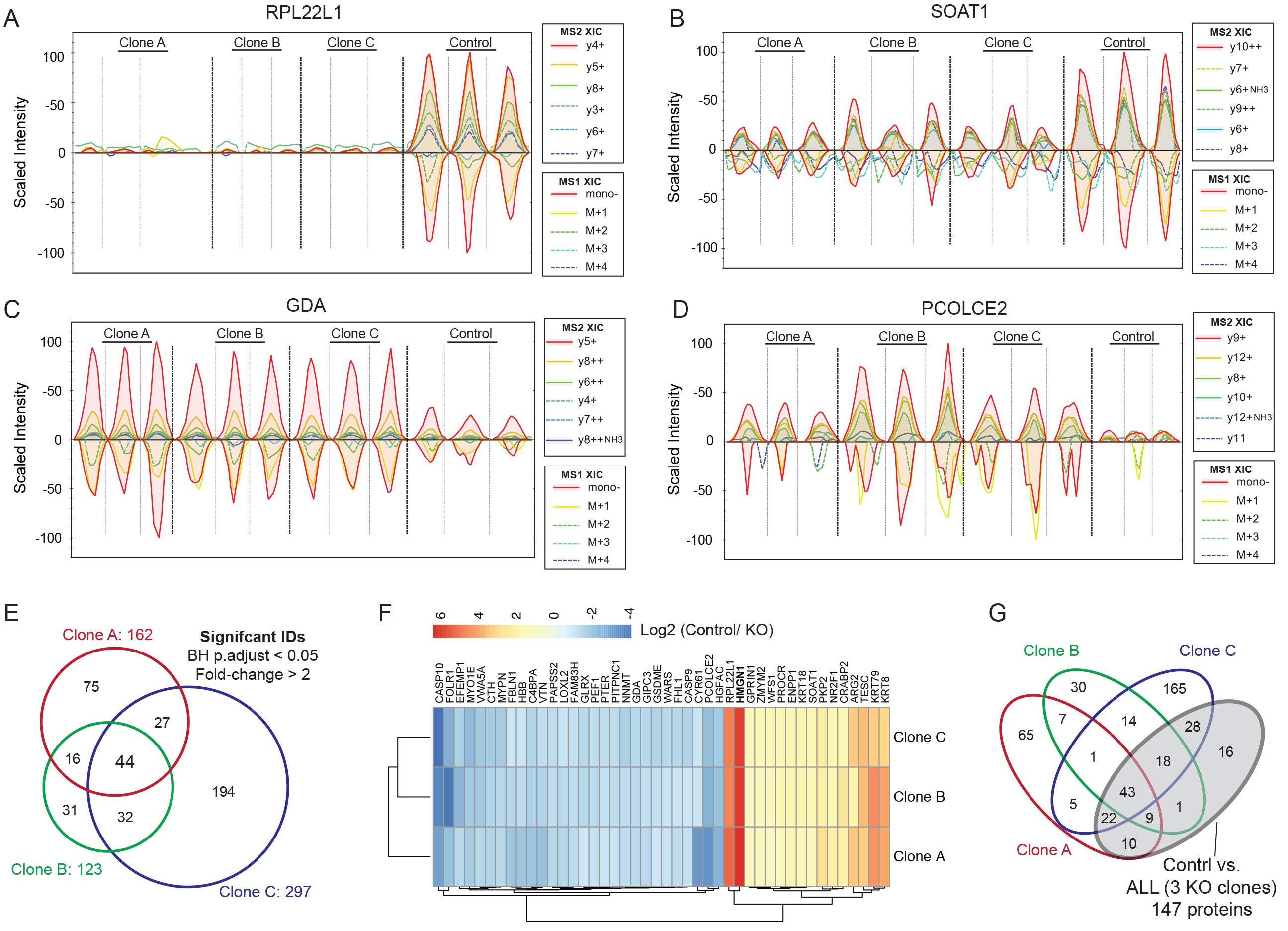
Reproducible and consistent DIA quantification on protein changes induced by HMGN1 knockout. **(A)** Extracted ion chromatography (XIC graphics) for TGNLGNVVHIER (Charge 3+) of RPL22L1 (Swiss-Prot ID: Q6P5R6) between 12 samples. **(B)** XIC graphics for NPTFLDYVRPR (Charge 2+) of SOAT1 (P35610). **(C)** XIC graphics for EWC[+57]FKPC[+57]EIR (Charge 3+) of GDA (Q9Y2T3). **(D)** XIC graphics for YC[+57]GDSPPAPIVSER (Charge 2+) of PCOLCE2 (Q9UKZ9). **(E)** Venn diagram of significant protein identities based on Clone A, B, and C. **(F)** Fold-changes of 44 overlapping, significantly regulated proteins in three clones. **(G)** Venn diagram of significant protein identities as (E). The grey circle denotes Control samples compared to Clone A, B, C combined as a knockout (KO) group, representing 147 proteins.

To uncover differentially expressed proteins between control and knockout clones, we performed statistical analysis visualized by volcano plots (**Figure 3**). A total of 162, 123, and 297 proteins were filtered as significantly regulated in Control vs. A, Control vs. B, and Control vs. C comparisons, by using BH adjusted P value <0.05 and fold-change >2 as the criteria.

Remarkably, HMGN1 was identified by six unique peptides and indeed was the protein that showed the largest fold-change in all of the three comparisons, with 83.42-, 67.14-, and 62.63- folds decreased from control to a quasi- noise level after CRISPR deletion. Of note, even in control samples, the HMGN1 protein intensity ranks below No. 3100, indicating that a less sensitive MS method quantifying e.g., ~3000 proteins will likely to fail in the precise quantification of HMGN1.

The quantification performance of DIA-MS on other differential proteins following HMGN1 depletion was visualized using extracted ion chromatographic peaks (XIC graphics, **Figure 4A-D** as examples). In such a plot, both the isotopic MS1 features for the peptide precursor (lower panel in XIC graphics) and the most intensive MS2 transitions whose number were specified during directDIA (upper panel in XIC graphics) could be aligned and compared across all samples. In the tryptic peptide examples representing for 60S ribosomal protein L22-like 1 (**Figure 4A,** protein level Control/CRISPR = 32.12 folds), Sterol O-acyltransferase 1 (**Figure 4B**, Control/CRISPR = 3.05 folds), Guanine deaminase (**Figure 4C**, Control/CRISPR = 0.365 fold), and Procollagen C-endopeptidase enhancer 2 (**Figure 4D**, Control/CRISPR = 0.150 fold), MS2- level quantification by DIA-MS was accurate and reproducible. A total of 44 differential proteins were profiled as significant in all the three comparisons, and harbored the consistent up- or down- regulations (**Figure 4E-F**). This indicates a reproducible proteomic perturbation after HMGN1 knockout. To benefit from the inclusion of *dish*- and *clone*- replicates, we then applied identical statistical criteria (BH adjusted P value <0.05 and fold-change >2) to the comparison between control (n=3) and all the knockout samples (n=9). Accordingly, 147 differential proteins were filtered for following up bioinformatic analysis.

To understand the biological consequences after HMGN1 deletion, we performed the enrichment analysis of GO and KEGG annotations on 147 proteins, taking the 6206 identified protein list as background (**Figure 5A**). We found that GO biological processes (BP) such as Cell differentiation (GO:0030154, P=7.52E-06), Regulation of cell proliferation (GO:0008285, P=0.0041), Receptor-mediated endocytosis (GO:0006898, P=1.07E-04), Negative regulation of endopeptidase activity (GO:001095, P=0.0092), and Proteolysis (GO:0006508, P=0.0292) were significantly enriched in the altered proteome, indicating the cell fate and endocytosis related processes were disturbed or remodeled. Many proteins annotated in “Negative regulation of cell proliferation” were found to be downregulated after HMGN1 deletion (**Figure 5A**), which is consistent to the mild but significant increase of cell proliferation observed in knockouts (**Supplementary Figure 3**). In agreement to the above BPs, cellular components (CC) such as Extracellular exosome (GO:0070062, P=4.79E-04), Integral component of plasma membrane (GO:0005887, P=0.0050), and Clathrin-coated vesicle (GO:0030136, P=0.0022) associated proteins were differentially regulated. Two KEGG pathways, Complement and coagulation cascades (P=5.72E-4) and ECM-receptor interaction (P=0.0343), were also enriched in the 147-protein list, reinforcing the newly discovered role of HMGN1 in immune response [33–36].

**Figure 5.**
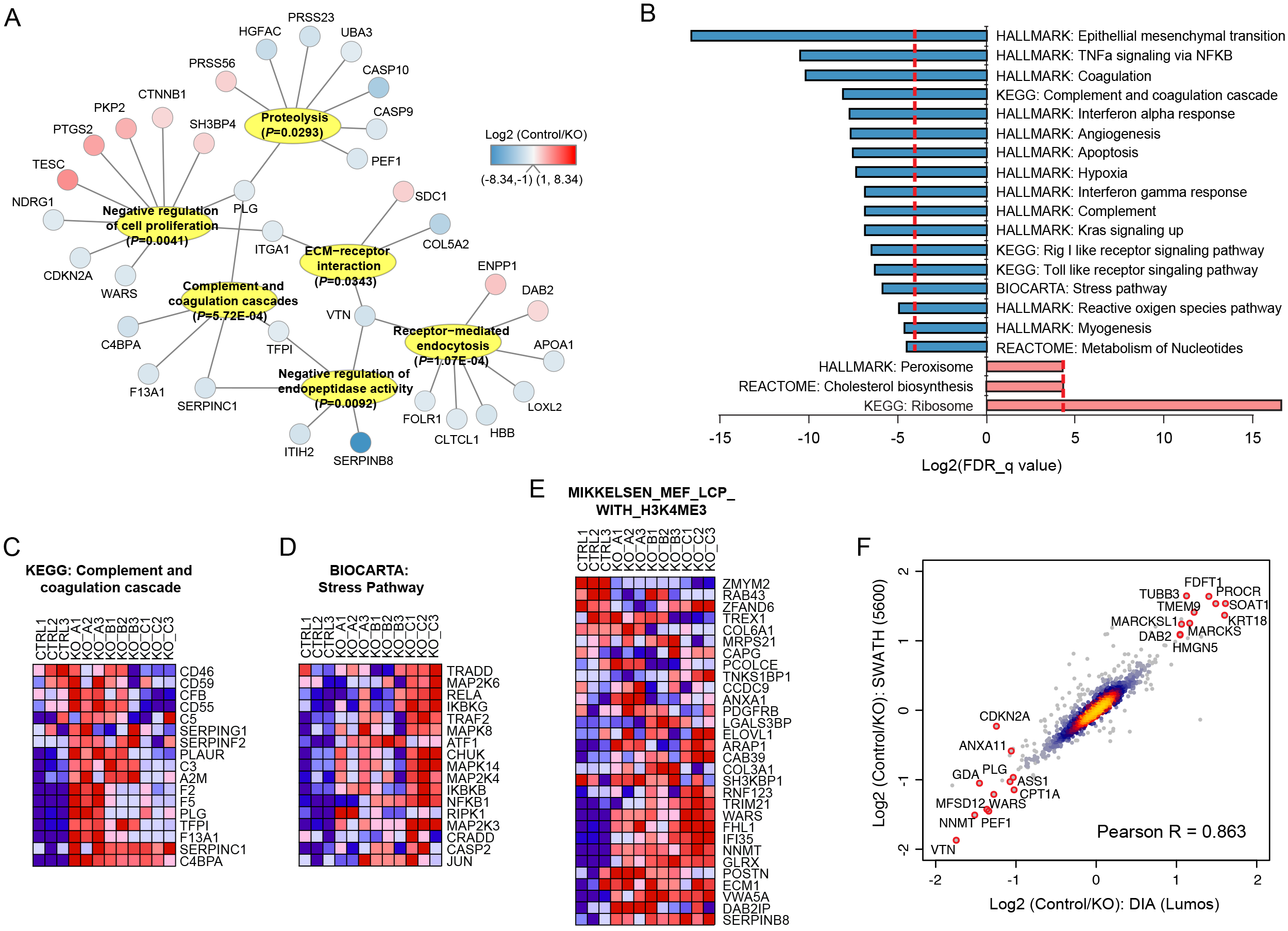
Annotation and enrichment analysis of proteome regulated after HMGN1 deletion. **(A)** Representative GO biological processes and KEGG pathways enriched in 147 protein list, and proteins annotated in each item. **(B)** GSEA analysis towards the full quantitative data set of all proteins using gene sets from KEGG, Reactome and HALLMARK summary. **(C-E)** Gene set examples enriched in KO group. **(F)** Foldchange correlation between SWATH-MS and DIA-MS based on directDIA approach.

To extend functional analysis to all the quantified proteins in a threshold-free manner, we applied gene set enrichment analysis (GSEA) in all HMGN1 deleted cells compared to wildtype controls. GSEA uncovered many biological regulations (**Figure 5B**, FDR *q*-values all below 0.05). In control samples, we found that the Peroxisome, Ribosome and Cholesterol biosynthesis functions were upregulated, suggesting these functions are impaired when HMGN1 is removed. In comparison, many more gene sets were enriched in knockout phenotypes, which could be summarized to a) Immune pathways, such as complement cascade and interferon alpha/ gamma responses, b) Cancer signaling, such as Rig I or Toll like receptor, TNFa, and Kras signaling, as well as angiogenesis, and c) Stress pathways, such as Reactive oxygen species pathway, hypoxia, and apoptosis. **Figure 5C** **and 5D** shows significant enrichment examples of Complement and coagulation cascades (KEGG) and Stress pathways (Biocarta) among upregulated processed after HMGN1 deletion. In particular, we found that among the H3K27-related datasets in the C2 CGP collection of MSigDB (Broad Institute) those genes marked by H3K27me3 (inactive histone marks) in reprogrammed iPS cells (MIKKELSEN_MCV6_HCP_ WITH_H3K27ME3) were enriched in the HMGN1 knockout condition (FDR =0.0021, **Figure 5E**), confirming the deletion of HMGN1 inactivates histone and chromatin activity [42]. Altogether, GSEA analysis helped in the further dissection of proteotypes induced by HMGN1 deletion.

Finally, as a global confirmation of the relative quantification results, we measured the same sample set by SWATH-MS method which is well established. Using panHuman library SWATH-MS profiled 3767 proteins (protein- FDR 1%). Indeed, the quantitative fold-changes derived from directDIA analysis in both SWATH and DIA were nicely correlated (Pearson R =0.863, **Figure 5F**), suggesting that both measurements accurately quantified protein regulations following HMGN1 deletion.

## Discussion

Our proteomic results shed the light on the cellular functions of HMGN1 protein. *Firstly*, as an abundant member, HMGN1 belongs to HMGNs, a family of proteins which binds specifically to the nucleosome core particle and reduces the compaction of the chromatin fiber [28]. HMGN1 has been discovered to modulate Histone H3 phosphorylation [28] and to remodel the chromatin during embryogenesis [43]. In this context, this protein was found to preferentially localize to DNase I hypersensitive sites, promoters, functional enhancers, and transcription factor binding sites [30] and has a variant-specific and tissue-specific effect on global transcription [31, 32]. Consistent to this major role of HMGN1 in regulating gene expression and DNA repair in mammalians [42], we found that those genes marked by H3K27me3 in reprogrammed iPS cells were enriched in knockout phenotypes using our proteomic data [44]. Even more interestingly, the same gene set was found to be enriched in wildtype cells when compared to HMGN1 overexpressed B cell progenitors, according to a recent transcriptome analysis [27]. Taking these results together, we can deduce that HMGN1 dosage, whether of overexpression, wildtype, and depletion, has a linear relationship with histone activity in regulating the overall gene expression.

*Secondly*, in previous reports, the Hmgn1^−/−^ mice were found to have reduced fertility, higher sensitivity to UV irradiation [29] and earlier signs of N-Nitrosodiethylamine-Induced liver tumorigenesis than their wildtype littermates [45]. Loss of HMGN variants affects the expression of stress-responsive genes in mouse embryonic fibroblasts (MEFs) [46]. We also found those stress pathways including hypoxia and apoptosis are induced after HMGN1 knockout, potentially explaining the vulnerability phenotypes discovered in Hmgn1^−/−^ mice.

*Thirdly*, the interaction of HMGN1 with chromatin was previously found to down-regulate the expression of N-cadherin, a transmembrane protein mediating cell-cell adhesion [47]. Notably, we detected significant proteomic changes (both up- and down- regulations) in the extracellular exosome and plasma membrane sub-proteomes, which might suggest that the “outside” proteome was already heavily rewired as a new state of HMGN1 deletion in the tested cell line. Further studies are therefore needed to delineate the relationship between HMGN1 downregulation and extracellular protein changes.

*Last but not the least*, HMGN1 recently was re-discovered to play an important role in immune response. It was identified as an alarmin, a group of mediators capable of promoting the recruitment and activation of antigen-presenting cells (APCs), including dendritic cells (DCs), that can potentially alert host defense against danger signals. Yang et al reported that extracellular HMGN1 acts as a novel alarmin that binds TLR4 and induces antigen-specific Th1 immune responses [33] and could be a potent immunoadjuvant by supporting T cell-mediated antitumor immunity [34, 35]. Recently a curative therapeutic vaccination regimen dubbed ‘TheraVac’ consisting of HMGN1 and resiquimod plus a checkpoint inhibitor was developed with the potential to eliminate various large tumors and induce tumor-specific immunity [36]. Our proteomic profiling is promising in this regard, because complement and coagulation cascades and interferon alpha or gamma responses were found to be significantly induced upon HMGN1 deletion, and therefore provides additional insights of HMGN1’s role in immunity.

The specific *clone*- and *dish*- effects we observed could be derived from possible CRISPR off-target effects, the real-time cell culturing difference, and/or experimental variations. There was no apparent cell proliferation difference between knockout clones. It is conceivable that proteotype provides closer scrutiny on phenotypes than mRNA profiling. Therefore, for a proteomic measurement on CRISPR effect, we recommend multiple *clone*- and *dish*- replicates to be included in the experiment to fully capture the proteotypic variations.

In summary, we applied a powerful DIA-MS measurement featuring decent sensitivity (>6200 proteins profiled), high reproducibility (R>0.98 between dish- replicates), high quantitative accuracy (for both absolute and relative scale quantification), cost-effectiveness (label-free quantification), and high sample throughput (2 hours per single measurement) in analyzing the cellular proteome after HMGN1 CRISPR deletion. Both the targeted protein (i.e., HMGN1) and the differential proteome closely associated to HMGN1 deletion were rapidly profiled. Our results provide new hints in understanding HMGN1 functions, and suggest the DIA-MS can be used as a rapid, robust, and reliable proteomic-screening step to assess the outcome of the CRISPR experiments.

## Methods

### Cell culture and transfection

T-REx-HeLa CCL2 cells were purchased from Invitrogen and cultured in DMEM (4.5 g/l glucose, 2 mM L-glutamin) (Gibco) supplemented with 10% fetal bovine serum (BioConcept), 100 U/ml penicillin (Gibco) and 100 μg/ml streptomycin (Gibco) at 37°C in a humidified incubator with 5% CO2. DNA transfection of the cells was performed with jetPrime (Polyplus) according to the manufactors’s instructions. The cell pellets were washed by pre-cold PBS twice and were snap frozen before proteomic sample preparation.

### MTT assay measuring cell proliferation

T-REx-HeLa CCL2 cells and CRISPR/Cas9-engineered HMGN1 knockout clones were seeded in 6-well plates and treated with 0.1 mg/ml MTT (3-[4,5-dimethylthiazol-2-yl]-2,5-diphenyltetrazolium bromide, Sigma) for 4h at 37°C. After removal of the media the converted dye was solubilized in 80% isopropanol, 20% DMSO. The absorbance of the converted dye was measured at a wavelength of 570 nm with background subtraction at 670 nm (Synergy HT, BioTek).

### CRISPR/Cas9-mediated gene knockout

CRISPR guideRNAs were designed based on their specificity score retrieved from the Optimized CRISPR Design web tool (http:crispr.mit.edu). Annealed DNA oligonucleotides containing the target sequence were cloned into the hSpCas9 plasmid (pX458, Addgene) using BbsI restriction sites. T-REx-HeLa cells were transfected with two hspCas9 constructs encoding gRNAs that target the second and third exon of the target gene HMGN1, respectively. The cell culture medium was replaced 4 hours after transfection and cells were recovered for 72 hours. For FACS sorting 1×10e6 cells were gently detached from the tissue culture plate with 0.25% trypsin-EDTA (Gibco) and resuspended in PBS containing 1% FBS. GFP-positive cells were isolated by FACS (BD Facs Aria IIIu sorter) and single cells were sorted into each well of a 96-well plate. Cell clones were expanded for three weeks and then screened for deletion events by western-blotting. Below lists the information of gRNA target sequences and oligonucleotides.

#### gRNA target sequences

gRNA_1_HMGN1_exon2: CCGCAGGTCAGCTCCGCCGAAGG

gRNA_2_HMGN1_exon3: CGCCGATCTCCTCTTGGGCTTGG

#### Oligonucleotides

Hmgn1_gRNA_1_exon2_forward: 5-CACCGCCGCAGGTCAGCTCCGCCGA-3

Hmgn1_gRNA_1_exon2_reverse: 5-AAACTCGGCGGAGCTGACCTGCGGC-3

Hmgn1_gRNA_2_exon3_forward: 5-CACCGCGCCGATCTCCTCTTGGGCT-3

Hmgn1_gRNA_2_exon3_reverse: 5-AAACAGCCCAAGAGGAGATCGGCGC-3

### Western-Blotting

For immunoblot analysis 1×10e6 T-REx-HeLa cells were lysed in 100 μl lysis buffer (0.5% NP40, 50 mM Tris-HCl, pH 8.0, 150 mM NaCl, 50 mM NaF supplemented with 1 mM PMSF and protease inhibitors (Sigma)). The cell lysate was cleared by centrifugation (15000xg, 20 min) and boiled for 5 min after addition of 3x Laemmli sample buffer. The denatured sample was loaded on NuPAGE 4-12% Bis-Tris SDS-PAGE gels (Invitrogen) for gel electrophoresis and then transferred onto nitrocellulose membranes (Trans-Blot Turbo, BioRad). Proteins were detected by rabbit polyclonal antibodies against HMGN1 (#5692, Cell Signaling) and mouse monoclonal antibodies against Actin (ab179467, Abcam). For immunodetection by enhanced chemiluminescence (ECL, Amersham) horseradish-peroxidase-coupled secondary antibodies (#7074, Cell Signaling and #115035003, Jackson ImmunoResearch) were used.

### Cell lysis and in-solution digestion

The protein extraction and digestion were preformed as previously described [23]. Briefly, cell pellets were suspended in 10M urea lysis buffer and complete protease inhibitor cocktail (Roche), ultrasonically lysed at 4°C for 2 minutes by two rounds using a VialTweeter device (Hielscher-Ultrasound Technology). The mixtures were centrifuged at 18,000 g for 1 hour to remove the insoluble material. The supernatant protein amount was quantified by Bio-Rad protein assay. Protein samples were reduced by 10mM Tris-(2-carboxyethyl)-phosphine for 1 hour at 37°C and 20 mM iodoacetamide in the dark for 45 minutes at room temperature. All the samples were further diluted by 1:6 (v/v) with 100 mM NH_4_HCO_3_ and were digested with sequencing-grade porcine trypsin (Promega) at a protease/protein ratio of 1:25 overnight at 37°C. The amount of the purified peptides was determined using Nanodrop ND-1000 (Thermo Scientific) and 1.5 μg peptides were injected in each LC-MS run.

### DIA mass spectrometry

The peptide separation was performed on EASY-nLC 1200 systems (Thermo Scientific) using a self-packed analytical PicoFrit column (New Objective) (75 μm × 60 cm length). This 50 cm column was packed with ReproSil-Pur 120A C18-Q 1.9 μm (Dr. Maisch GmbH). An LC method of 2 hours was used (Buffer A: 0.1% formic acid, buffer B: 80% acetonitrile containing 0.1% formic acid). Briefly, a 5% to 37% Buffer B gradient of 109-min was conducted with the flow rate of 300 nl/min at 60 °C in a column oven (PRSO-V1, Sonation GmbH).

The Orbitrap Fusion Lumos Tribrid mass spectrometer (Thermo Scientific) with a NanoFlex ion source was coupled to the LC platform. Spray voltage was set to be 2,000 V and heating capillary was kept at 275 °C. The DIA-MS method consisted of a MS1 survey scan and 40 MS2 scans of variable windows. The MS1 scan range is 350 - 1650 m/z and the MS1 resolution is 120k at m/z 200. The MS1 full scan AGC target value was set to be 2.0E5 and the maximum injection time was 100 ms. The MS2 resolution was set to 30,000 at m/z 200 and normalized HCD collision energy was 28%. The MS2 AGC was set to be 5.0E5 and the maximum injection time was 50 ms. The default peptide charge state was set to 2. Both of MS1 and MS2 spectra were recorded in profile mode.

The additional SWATH-MS measurements were performed as previously published [26].

### DIA data extraction and analysis

The DIA runs were analysed with Spectronaut Pulsar version 12.0 (Biognosys AG, Switzerland) [38, 48]. In the directDIA approach, DIA runs were directly searched against Swiss-Prot protein database (March 2018, 20,258 entries) with following settings: full tryptic allowing two missed cleavages, set carbamidomethylation as a fixed modification on all cysteines, set oxidation of methionines, protein N-terminal acetylation as dynamic modifications. Both precursor and protein FDR were controlled at 1%. As for quantification, interference correction function was enabled, and top 3 peptide precursors were summed for protein quantification. Data filtering was done with Qvalue sparse in this 12-sample experiment. In panHuman approach, the published set of mass spectrometric assays for 10,000 human proteins was downloaded [40] and the protein quantitative results were summarized in the same way as directDIA. In both approaches, all the other parameters in Spectronaut are kept as default unless mentioned. To combine directDIA and panHuman results, the protein quantification results of those only acquired by panHuman library (but not by directDIA) were simply appended into the directDIA results.

### Bioinformatic analysis

Hierarchical clustering analysis (HCA) was performed by Cluster 3.0 on the normalized, log2-transformed, 2-dimensional-centered DIA-MS intensities and visualized by TreeView. Principal component analysis (PCA) was performed by R software on the normalized, log2-transformed data. Quantitative CVs were calculated by using normalized DIA data at the original scale. For differential expression analysis, P values were corrected for multiple testing with the Benjamini-Hochberg (BH) method [49]. DAVID Bioinformatics Resources 6.8 (https://david.ncifcrf.gov/) [50] was used to perform the enrichment analysis of biological processes, cellular component and pathways in the 147-protein list. Representative proteins and their enriched items were visualized by Cytoscape v3.70 [51]. GSEA 3.0 (Gene set enrichment analysis [44]) was downloaded from http://software.broadinstitute.org/gsea/index.jsp, and default settings were used to infer the degree to which a given gene set is overrepresented at the upper or lower ends of a ranked list of genes by at an estimated probability (i.e., FDR-q value). Briefly, a running-sum statistic is generated from the ranked list of genes, with the magnitude of the increment depending on the correlation of the gene with the phenotype (control and HMGN1 deletion).

## Supporting information

## Acknowledgement

We thank Lukas Reiter and Roland Bruderer from Biognosys AG for the technical assistance. We would like to particularly thank Dr. Ruedi Aebersold for the resource support and helpful discussions. M.M. was supported by a Long-Term Fellowship from the European Molecular Biology Organization (ALTF 928-2014).

