## Supplementary material for "Rapid Proteomic Screen of CRISPR Experiment Outcome by Data Independent Acquisition Mass Spectrometry: A Case Study for HMGN1"

1. Department of Biology, Institute of Molecular Systems Biology, ETH Zurich, Zurich, Switzerland
2. Yale Cancer Biology Institute, Yale University, West Haven, CT 06516, USA
3. Department of Genome Integrity, Institute of Molecular Genetics of the Czech Academy of Sciences, Prague, Czech Republic
4. Department of Pharmacology, Yale University School of Medicine, New Haven, CT 06520, USA
5. These two authors contribute equally to the study.

### Supplementary Figure 1

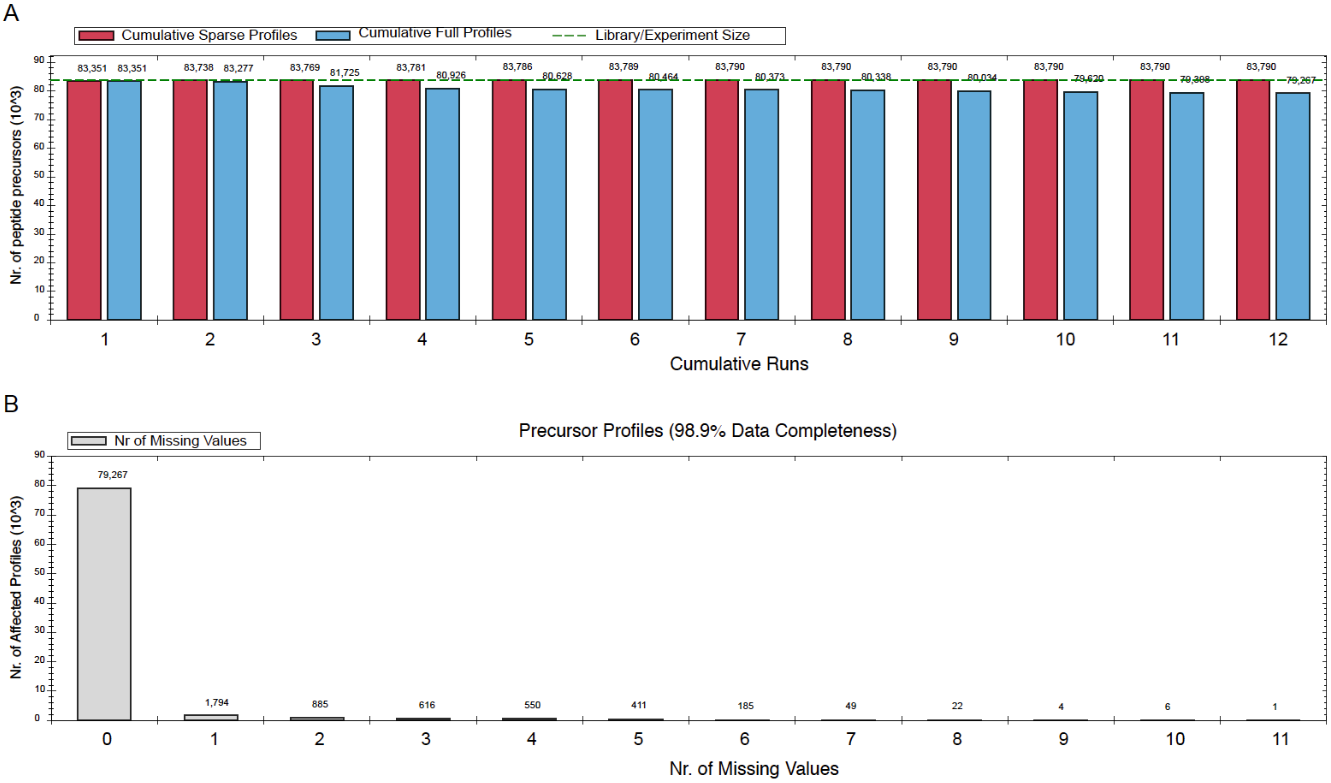


**Figure S1. Data recovery and completeness by directDIA analysis.** **(A)** Cumulative of peptide precursor identifications across 12 LC-MS runs. **(B)** Distribution of missing values. Note that nearly 80,000 precursors were identified in every sample.

**Supplementary Figure 2**

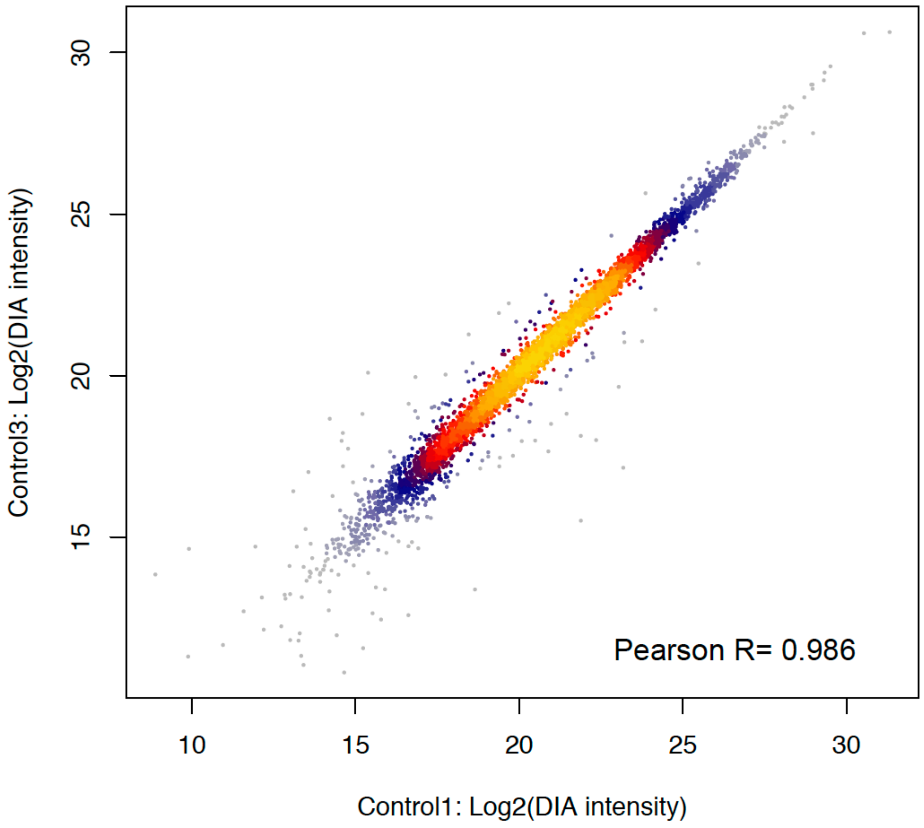


**Figure S2. Example of quantitative correlation of DIA-MS intensity between two control samples (*dish-* replicates).** For the summary of Pearson correlation coefficients between all the samples, please refer to Figure 2D in the main text.

**Supplementary Figure 3**

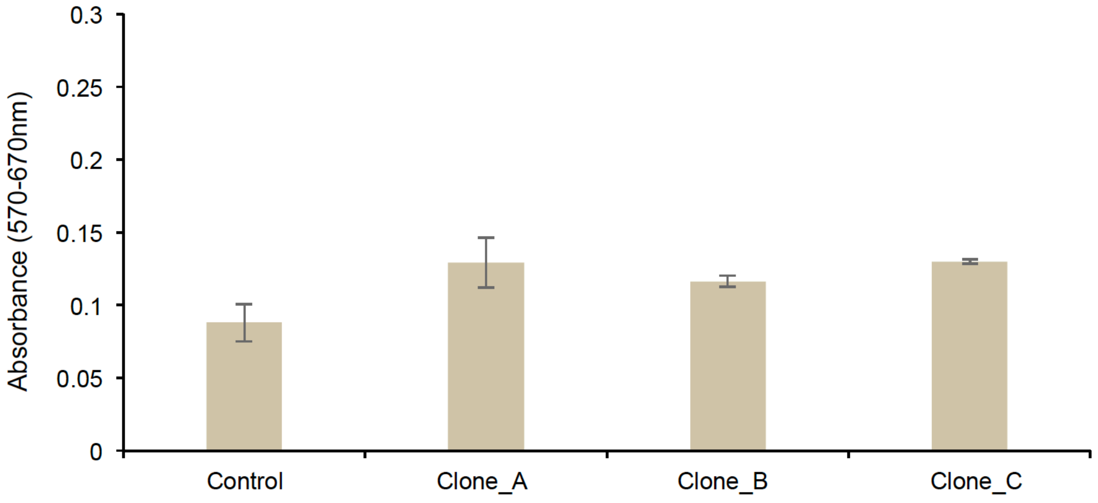


**Figure S3. MTT assay assessing the cell proliferation of different clones.** Note there are no statistical difference between three knockout clones (A, B, and C). However, there are mild increases of cell proliferation rate in all knockout clones when compared to controls ($P<0.05$). Error bars represent 95% confidence interval.

**Supplementary Figure 4**

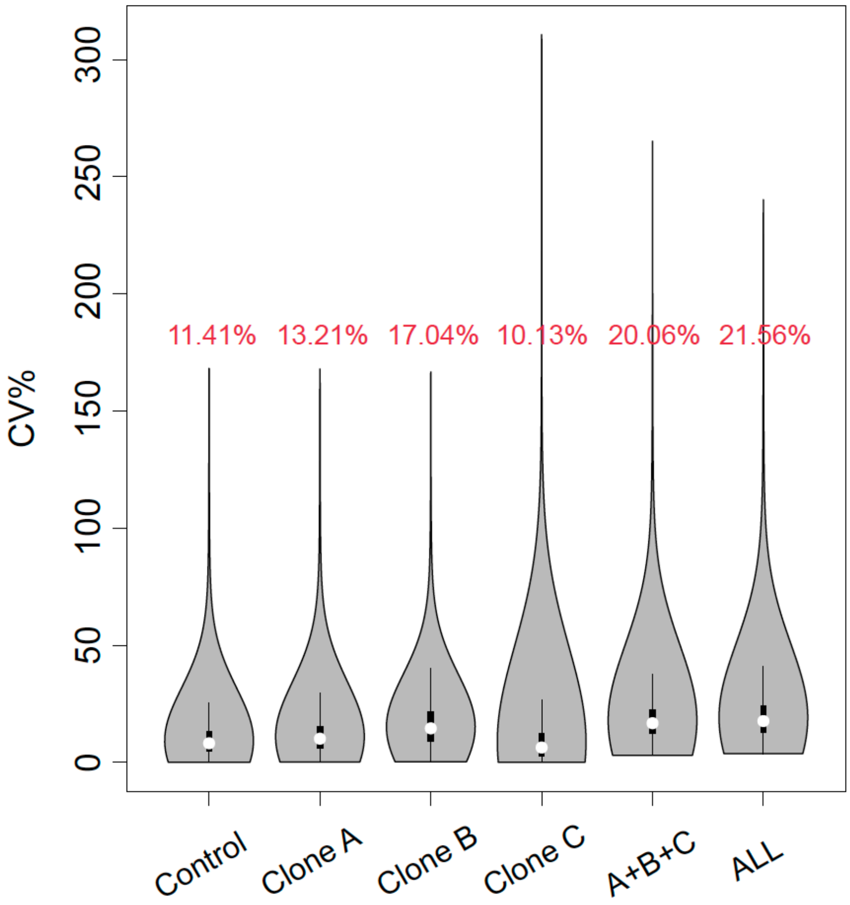


**Figure S4. CV distribution for dish- replicates in each clone.** Note violin plot of CV calculated based on dish- replicates for each of the control, HMGN1 deletion clones, all knockout clones combined, and all the samples combined is shown. The red numbers denote the average CV.
